# Post-daunorubicin treatment effects on cardiovascular function in the Ts65Dn mouse model of Down syndrome

**DOI:** 10.64898/2026.05.21.726916

**Authors:** Michelle A. Buckman, Anastasiia Vasileva, Hardik Kalra, Immaculate Edwin, Guiru Ma, Mikhail Vasilyev, Charles R. Jedlicka, Barry London, Gary Beasley, Patrick Breheny, Michael H. Tomasson, Melissa L. Bates

## Abstract

Adults with Down syndrome (DS) are two times more likely to be diagnosed with chronic heart failure post-anthracycline chemotherapy compared to age and sex-matched adults without DS. They have an elevated lifetime risk of cardiovascular diseases, increasing their likelihood of anthracycline-induced chronic cardiovascular toxicity. We investigated the chronic effects of daunorubicin on the cardiovascular system of the adult Ts65Dn mouse model of DS compared to wild type euploid mice (WT). WT and Ts65Dn mice received two doses of 2mg/kg or 4mg/kg of daunorubicin or saline and were monitored for up to 117 days. Cardiac and vascular function were evaluated using left ventricular catheterization, histology, pulse wave velocity, and cardiac troponin tests. Survival significantly decreased in the Ts65Dn 4mg/kg group compared to saline controls (p<0.001). Further experiments were carried out with the saline and 2mg/kg groups, which exhibited lower mortality, more consistent with chronic toxicity. Body weight (p=0.001), end-diastolic pressure (p=0.016), and left ventricular mass (p=0.021) decreased in treated mice. The effect of treatment differed significantly between strains for ejection fraction (p=0.029). Pulse wave velocity increased over time (p<0.001). A significant interaction between treatment and strain was observed for collagen in both the left ventricles and thoracic aorta (p=0.002 and p<0.001, respectively). There was a strain difference for cardiac troponin I, indicating an increase in Ts65Dn mice (p=0.020). Daunorubicin treatment results in a distinct cardiovascular remodeling phenotype in Ts65Dn mice. More mechanistic studies are warranted to outline the pathophysiology of anthracycline cardiovascular toxicity in DS.

## INTRODUCTION

The population prevalence of DS, most commonly caused by the triplication of human chromosome 21, has increased four-fold since 1950, resulting in a rate of 6.7 cases per 10,000 inhabitants (1). Additionally, there has been a 3.75-fold increase in the average life expectancy of people with DS since 1970 (2). While there are well-established health supervision guidelines for children with DS, consensus-based guidelines for adults with DS are lacking, despite the increase in their life expectancy (3).

We previously found that adults with DS are three times more likely to be diagnosed with malignancies of the lymphoid, hematopoietic, and related tissues, including leukemia and myeloma, compared to age and sex-matched controls without DS (4). They also have a lower incidence of breast, cervical, lung, and prostate cancer and a similar incidence of endometrial and colon cancer (4). Survivorship outcomes, including risk of chronic cardiovascular disease following chemotherapy, are not well quantified in the DS population.

Anthracyclines are first-line chemotherapy agents for many cancers, including leukemia, lymphoma, and breast cancer (5). Chronic cardiovascular dysfunction is observed in survivors of cancer after anthracycline chemotherapy, and children with DS are at an increased risk of chronic cardiotoxicity after anthracycline treatment (5, 6). Similarly, we previously found that individuals with DS, treated with anthracyclines as adults, are two times more likely to be diagnosed with heart failure, a classic manifestation of anthracycline cardiotoxicity, compared to age- and sex matched controls without DS (4). Elevated chronic cardiotoxicity risk may be exacerbated by increased pre-existing cardiovascular disease risk or different mechanisms of cardiovascular injury in the DS population (7). However, it is unclear if this increased risk is an innate physiological response due to the 1.5-fold increase in gene dosage on human chromosome 21, pre-existing co-morbidities, or differences in lifestyle or healthcare access that may increase cardiovascular disease risk. This emphasizes the need for further studies on the pathophysiology of anthracycline cardiotoxicity in a model of DS without confounding lifestyle factors or pre-existing co-morbidities.

We leveraged a mouse model of DS with increased gene dosage of human chromosome 21-related genes to better understand the pathophysiology of anthracycline cardiotoxicity in DS. The Ts65Dn mouse model is the most widely studied mouse model of DS, and is characterized by segmental trisomy at the distal end of mouse chromosome 16, a region that shares conserved synteny with human chromosome 21 and the proximal end of mouse chromosome 17 (8, 9). The Ts65Dn mouse model has been used to investigate cardiovascular phenotypes of DS, such as blood pressure, heart rate variability, and congenital heart defects (10–12). We hypothesized that daunorubicin treatment would enhance the risk of chronic cardiovascular toxicity after cessation of treatment in the Ts65Dn mouse model of DS.

## MATERIALS AND METHODS

### Animals

Four-month-old Ts(17<16>)65Dn (Ts65Dn) mice and control wildtype Ts(17<16>)65Dn (WT) mice (Strain No. 005252; B6EiC3Sn.BLiA-Ts (17^16^)65Dn/DnJ) were obtained from the Jackson Laboratories. Mice were housed in the animal facility at the University of Iowa in standard housing conditions, including an ambient temperature of 20°C to 26°C, a 12:12h light-dark cycle, and ad libitum access to standard rodent chow (NIH-31 Rodent Diet, Envigo) and water. All procedures were approved by the University of Iowa Institutional Animal Care and Use Committee.

### Experimental design

First, a dose-finding study was performed to obtain a dose with minimal acute lethality to assess the chronic effects of anthracyclines, but not acute cardiotoxicity. In the absence of existing protocols for evaluating daunorubicin cardiotoxicity in Ts65Dn mice, we adapted a protocol previously used for C57BL/6J mice, where the authors investigated vascular function after doxorubicin treatment, a drug in the same class as daunorubicin (13). [Figure 1]. Mice were divided into three groups based on the daunorubicin concentration: 1) 2mg/kg treatment group, 2) 4mg/kg treatment group, and 3) saline control group. Each group consisted of four male and four female WT mice (total = 24 mice) and six male and six female Ts65Dn mice (total = 36 mice). Mice were intraperitoneally injected with daunorubicin or saline on days 0 and 7, and survival was monitored for up to 117 days. (Figure 1).

**Figure 1.**
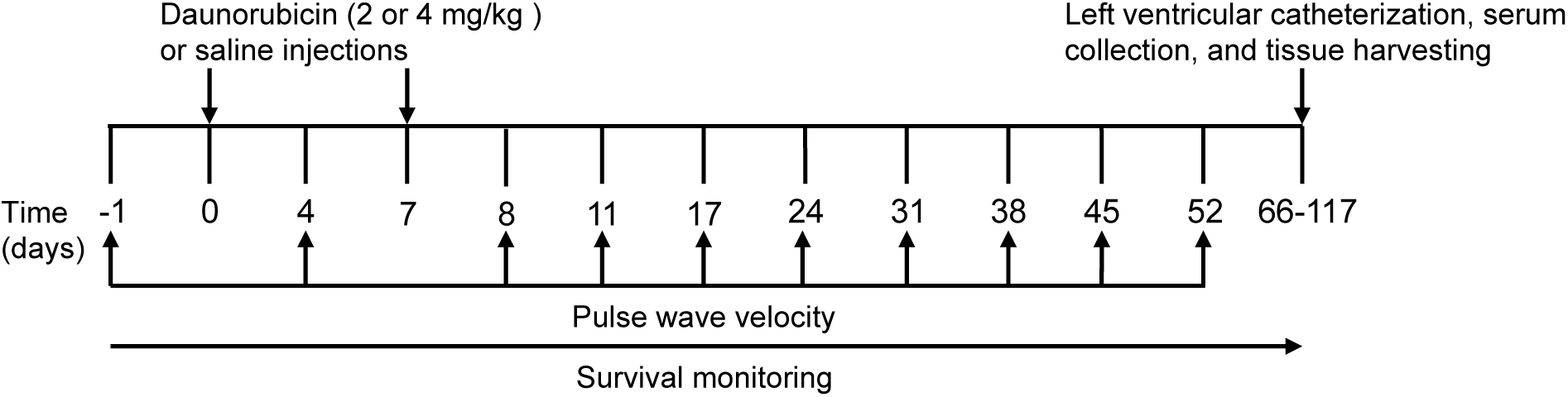
Overall experiment timeline. Four-month-old male and female WT (n=8/group, n=24 total) and Ts65Dn (n=12/group, n=36 total) mice were randomly divided into three groups according to the dose of daunorubicin (2 mg/kg and 4 mg/kg), with saline as a control. Abdominal aorta pulse wave velocity was assessed at multiple time points, followed by left ventricular catheterization, and serum and tissue collection.

### Pulse wave velocity

Pulse wave velocity was measured non-invasively using the Indus Doppler Flow Velocity System (Indus Instruments, Webster, TX) before the start of daunorubicin treatment (day -1) and subsequently on days 4, 8, 11, 17, 24, and was continued once a week for 53 days. [Figure 1]. Abdominal-aortic pulse wave velocity was measured as previously described (13). Anesthesia was induced with 3-4% isoflurane. Mice were maintained on a 36°C heated platform, with their snout in a nose cone and anesthesia maintained at 1-1.5%. Flow velocity signals were measured simultaneously with a 10-MHz and a 20-MHz probe. The 10-MHz probe was positioned on the left of the upper mid chest to obtain the aortic arch velocity signal. The 20-MHz probe was placed on the mid-lower abdomen, perpendicular to the aorta, to obtain the abdominal velocity signal. Micro-positioners were used to support the probes, and the distance between both probes was measured using calipers. The Doppler Signal Processing Workstation (Indus Instruments, Webster, TX) was used to analyze spectrograms to obtain the transit time. Pulse wave velocity was calculated as the distance between the probes (in centimeters) divided by the transit time (in seconds).

### Left ventricular catheterization

Left ventricular catheterization was performed as previously described (14). Mice were anesthetized with intraperitoneal injections of urethane (Sigma-Aldrich) dissolved in sterile 0.9% NaCl (Hospira, Inc.). Each mouse received an initial dose of 1.25 g/kg with two additional doses of 0.75 g/kg and 0.5 g/kg as needed to achieve a surgical plane of anesthesia, confirmed by the absence of toe pinch and palpebral reflexes (15). Before data collection, a single-segment pressure-volume catheter (PVR1030; ADInstruments) was calibrated using the Millar Mikro-Tip Pressure Volume System (ADInstruments, Oxford, UK) following 30 minutes of soaking in prewarmed saline. The calibrated catheter was inserted into the right common carotid artery, through the aorta, into the left ventricle. Instantaneous steady-state measurements of pressure and volume were recorded for a minimum of 5 minutes at a sampling rate of 1000 Hz using the Lab Chart 8.0 software package (AD Instruments, Colorado Springs, CO). Cardiac output, stroke volume, and ejection fraction were calculated as we have previously described (16). Consistent with our prior work, heart rate variability analyses were also performed using the Lab Chart 8.0 software (15). Mice were euthanized at the end of the experiments by harvesting vital tissues.

### Histology

The thoracic aorta and left ventricle were immediately harvested after left ventricular catheterization. The left ventricle with the septum and the right ventricle were weighed to calculate the Fulton index as the ratio of the right ventricle and the left ventricle with the septum, (Right ventricle/Left ventricle + Septum) (17). The tissues were fixed in formalin overnight and stored at 4°C in 70% NaOH for up to four months. The tissues were embedded in paraffin and serially sectioned at 5 µm. The left ventricle was stained with Masson’s trichrome collagen stain as a measure of cardiac fibrosis. The thoracic aorta was stained with picrosirius red for collagen and Verhoeff-Van Gieson for elastin. All images were captured using an Olympus BX-63 brightfield microscope at a magnification of 20x with the CellSens software at the University of Iowa’s Central Microscopy facility. An average of ten sections per mouse was used to quantify the blue-stained collagen-positive areas of the left ventricle, and six sections for the red-stained collagen and black-stained elastin of the thoracic aorta (ImageJ, National Institutes of Health and the Laboratory for Optical and Computational Instrumentation, University of Wisconsin). Regions of interest with an area of 500,000 µm^2^ were created on each section of the left ventricle with the grid option of ImageJ. Within each section, up to 20 regions of interest were randomly selected using a random number generator in Microsoft Excel. Fibrosis was quantified as the area occupied by collagen divided by the total area and expressed as a percentage for each region of interest. The thickness of individual smooth muscle sheets and the overall length of the tunica media of the thoracic aorta were recorded to evaluate vessel morphology. The percentage area of collagen (red) and elastin (black) in the tunica media of the thoracic aorta was manually segmented and quantified. To ensure unbiased quantification, the analysis was performed by two blinded investigators.

### Enzyme-linked Immunosorbent Assay (ELISA)

Blood was collected from the mice by cardiac puncture after left ventricular catheterization, and serum was stored at -80°C. Cardiac troponin I was quantified using the high-sensitivity mouse cardiac troponin-I ELISA kit (Life Diagnostics, Inc., Cat. No. CTNI-1-HS) as per the manufacturer’s instructions.

### Statistical analysis

Overall survival was evaluated using the Kaplan-Meier statistical test in GraphPad Prism 10.0. Pairwise comparisons of individual survival curves were used to detect significance using the Log-rank (Mantel-Cox) test. Bonferroni corrections were used for multiple comparisons of survival curves comparing each treatment group with the saline control group for each strain (p<0.0125. All graphs were generated using the GraphPad Prism 10.0 software package. All other statistical analyses were conducted using the Minitab statistical software package (State College, PA) with a significance level of p<0.05 and a 95% confidence interval. Analysis of variance (ANOVA) was used to evaluate differences in hemodynamic and histological parameters with strain and treatment as fixed variables, and strain × treatment interaction included in the model. Analysis of covariance (ANCOVA) was used for pulse wave velocity and histology analyses, with days and number of sections as continuous variables, respectively.

## RESULTS

### Survival

Survival significantly decreased in Ts65Dn mice treated with 4mg/kg of daunorubicin compared to the Ts65Dn saline group (p<0.001). (Figure 2). Importantly, there was no significant difference in the overall survival between the saline groups and the 2mg/kg group for both strains. (Supplemental Table S1). The 4 mg/kg group was excluded from further analysis, as this dose resulted in a significantly higher acute mortality rate. All other experiments were continued with the 2mg/kg groups.

**Figure 2.**
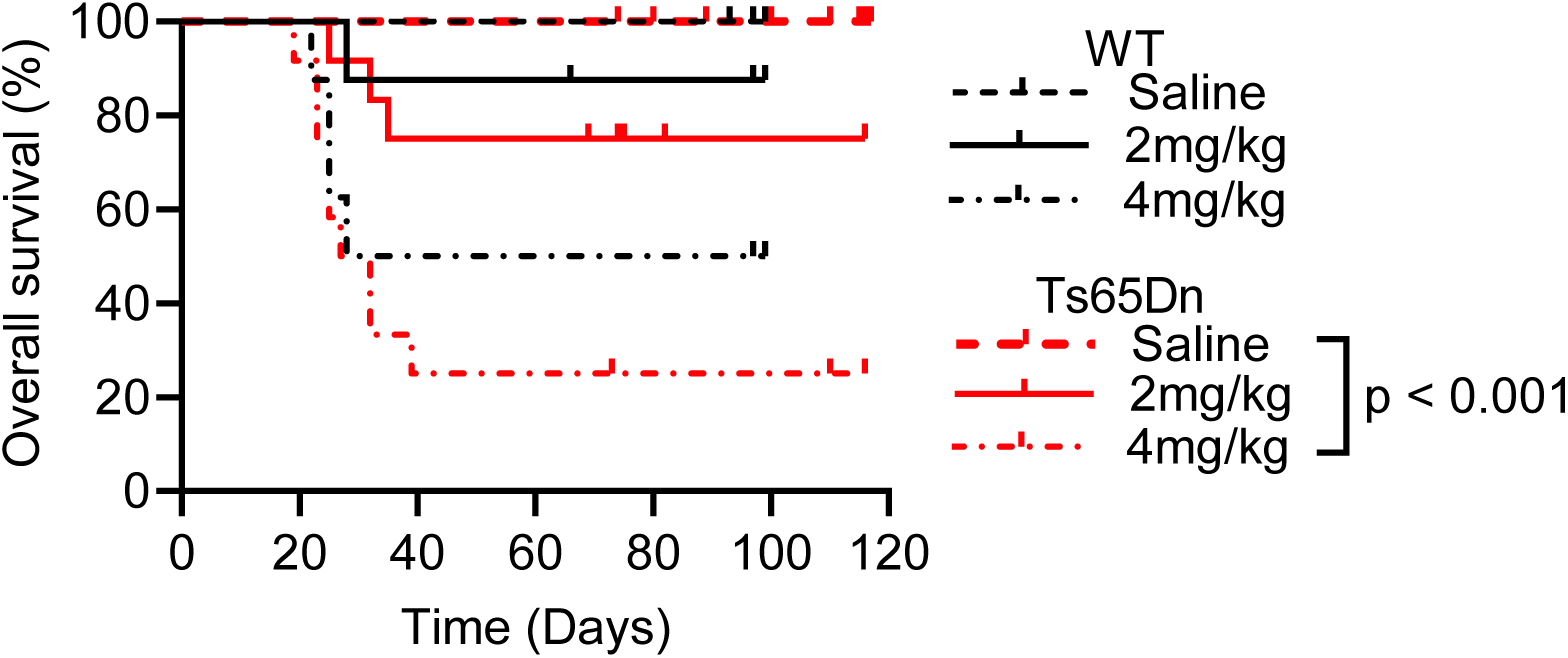
Overall survival. Higher dose (4mg/kg) of daunorubicin reduced survival in Ts65Dn mice. Pairwise comparisons of individual survival curves were used to detect significance after correcting for multiple comparisons. Survival of Ts65Dn mice treated with 4mg/kg of daunorubicin was significantly lower compared to the saline group (p < 0.001).

### Cardiac function

The body weight of mice at the end of the experiment and end-diastolic pressure significantly decreased in both treated groups (p=0.001 and p=0.016, respectively). (Table 1). A significant interaction between strain and treatment was observed for ejection fraction (p=0.029), consistent with an observed increase in WT-treated mice and a decrease in Ts65Dn-treated mice. There was no significant change in end diastolic volume, end systolic volume, heart rate, stroke volume, and cardiac output. (Table 1). Left ventricular mass significantly decreased in treated mice (treatment, p=0.021). There was no significant difference in heart rate variability between groups (Supplemental Table S2). Since there was no significant difference in heart rate between groups, we did not perform M-curve analyses of heart rate variability as we did in our previous study (15).

**Table 1.**
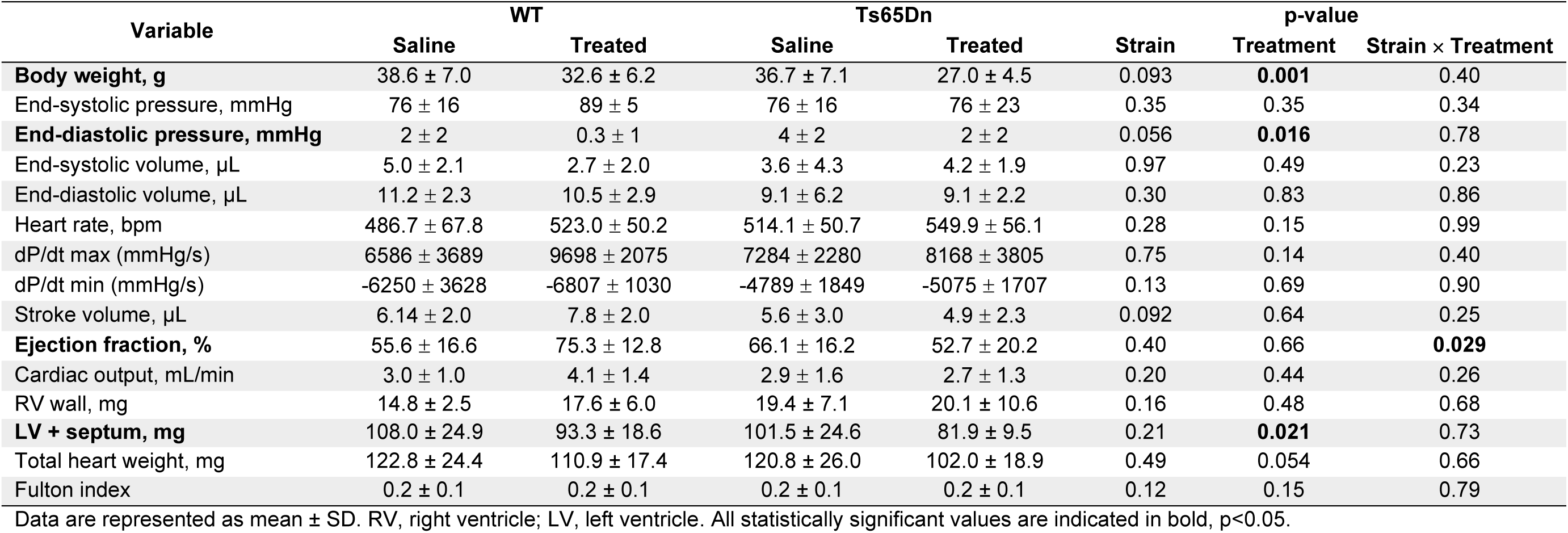
Summary statistics of baseline cardiac function and ventricular morphology.

Quantification of the percent area of the left ventricle containing collagen revealed a significant strain × treatment interaction (p<0.001), consistent with increased fibrosis in WT treated mice and decreased fibrosis in Ts65Dn treated mice (Figure 3).

**Figure 3.**
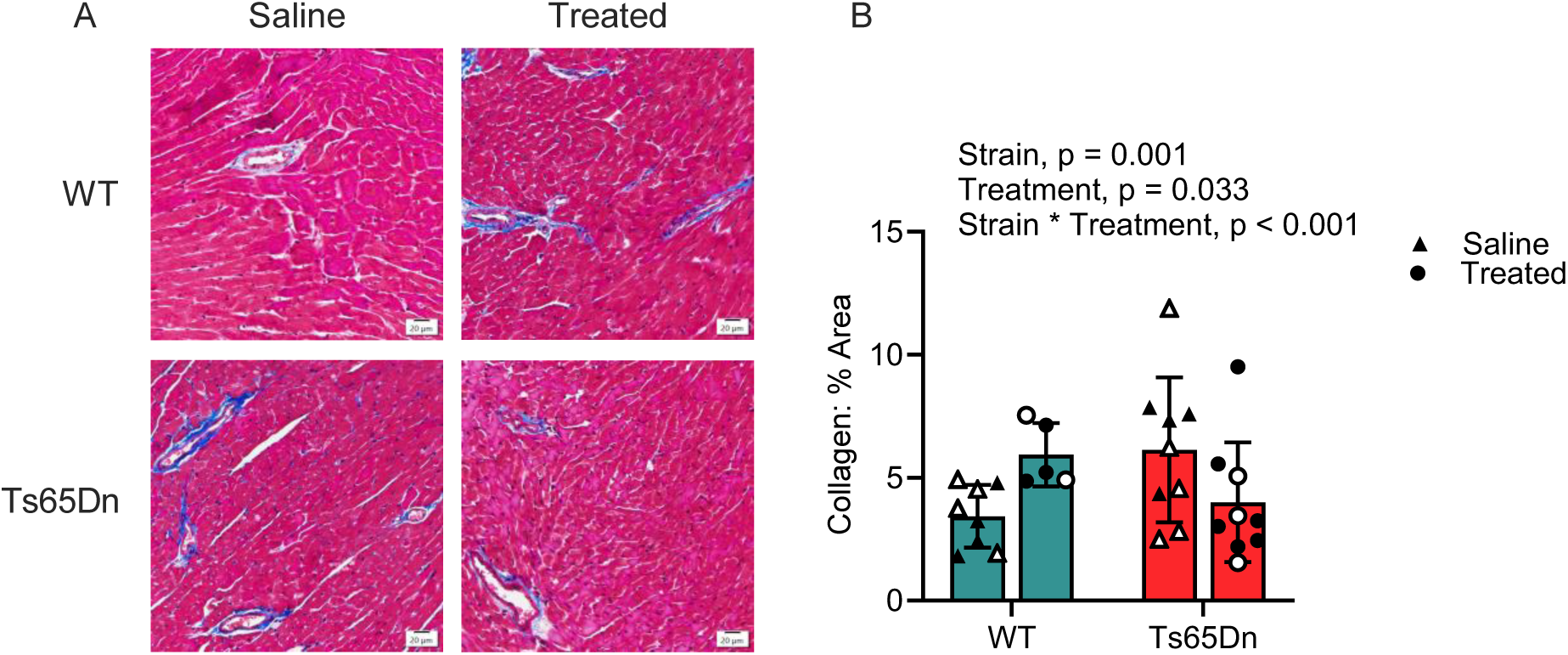
Cardiac morphology. Cardiac fibrosis decreased in the left ventricles of Ts65Dn-treated mice. (A) Representative images showing Masson’s trichrome staining of the left ventricles. Images were captured at a magnification of 40x with a scale bar of 50μm. (B). Quantification of the % area of the left ventricles occupied by collagen (strain, p = 0.001, treatment, p = 0.033, strain × treatment, p < 0.001). Closed shapes represent female mice, and open shapes represent male mice.

### Vascular function

Pulse wave velocity increased with time similarly in both strains and treatment groups (p<0.001). (Figure 4A and Supplemental Table S3). We found a significant treatment and strain × treatment effect for the thickness of individual smooth muscle sheets of the thoracic aorta (p<0.001 and p=0.023, respectively). (Figure 4B). Similarly, a significant treatment and strain × treatment effect was observed for the thickness of the tunica media of the thoracic aorta (p<0.001 and p=0.003, respectively). (Figure 4C). Interestingly, we found a significant interaction between strain and treatment for collagen (p=0.002), showing an increase in collagen in the thoracic aorta of WT treated mice and a decrease in collagen in the thoracic aorta of Ts65Dn treated mice. (Figure 4D). However, there was no significant effect of strain, treatment, or strain × treatment on elastin in the thoracic aorta. (Figure 4E).

**Figure 4.**
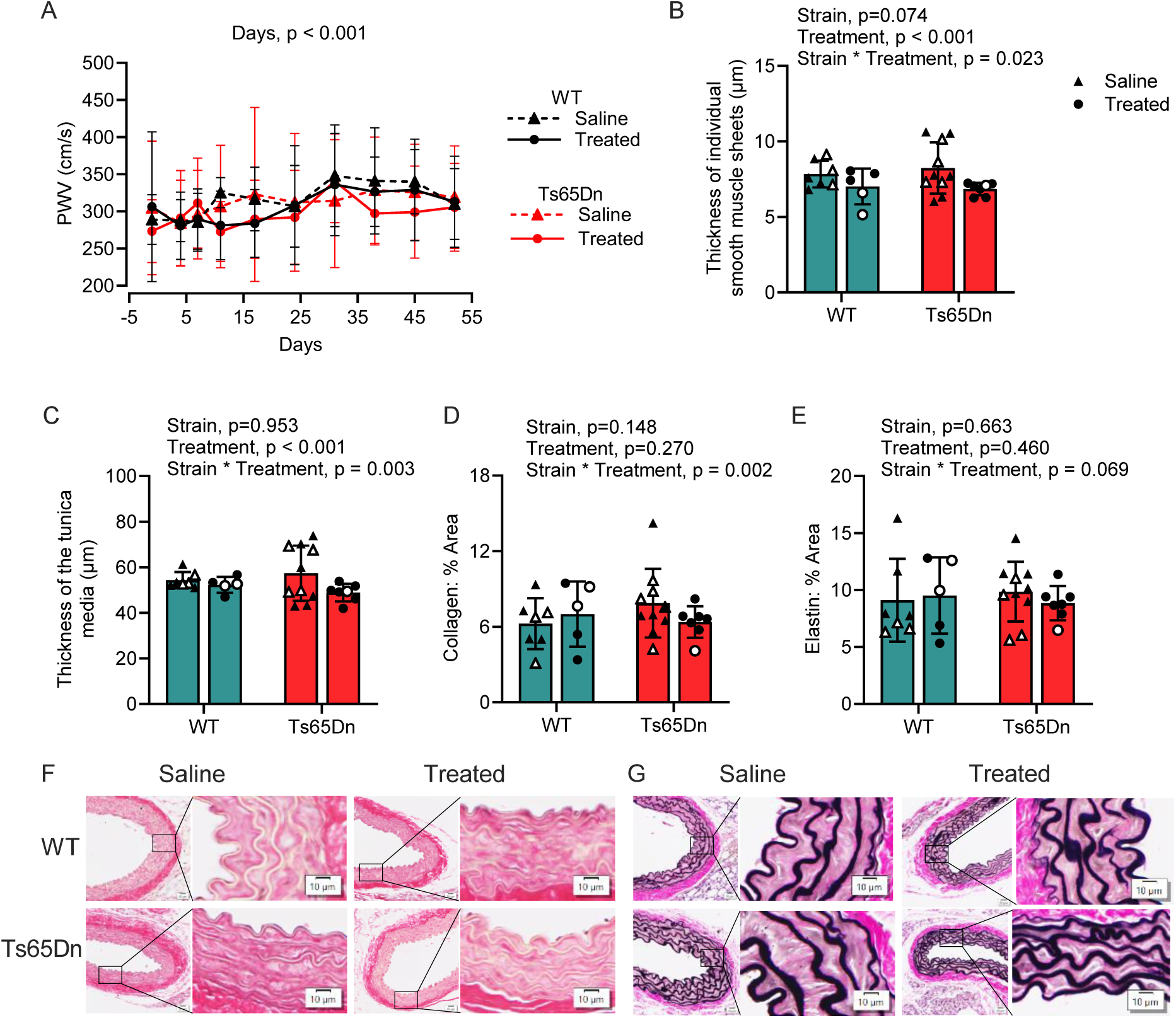
Arterial stiffness and morphology. Daunorubicin treatment altered vascular structure in Ts65Dn mice. (A). Abdominal aorta pulse wave velocity measurements. Pulse wave velocity significantly increased with time (days, p < 0.001). (B). Thickness of individual smooth muscle sheets in the tunica media of the thoracic aorta (treatment, p <0.001 and strain × treatment, p = 0.003). (C). Overall thickness of the tunica media of the thoracic aorta (treatment, p < 0.001 and strain × treatment, p = 0.003). (D). Quantification of collagen in the thoracic aorta (strain × treatment, p = 0.002). (E). Quantification of elastin in the thoracic aorta. (F). Representative images showing Picro-Sirus red collagen staining and (G). Verhoeff-van Gieson staining for elastin in the thoracic aorta. All data are represented as mean ± SD. Images were captured at a magnification of 40x with a scale bar of 20μm. Closed shapes represent female mice, and open shapes represent male mice.

### Troponin

Ts65Dn mice had elevated concentrations of high-sensitivity cardiac troponin I (p=0.020). There was no significant effect of treatment or strain × treatment (p=0.240 and p=0.862, respectively). (Figure 5).

**Figure 5.**
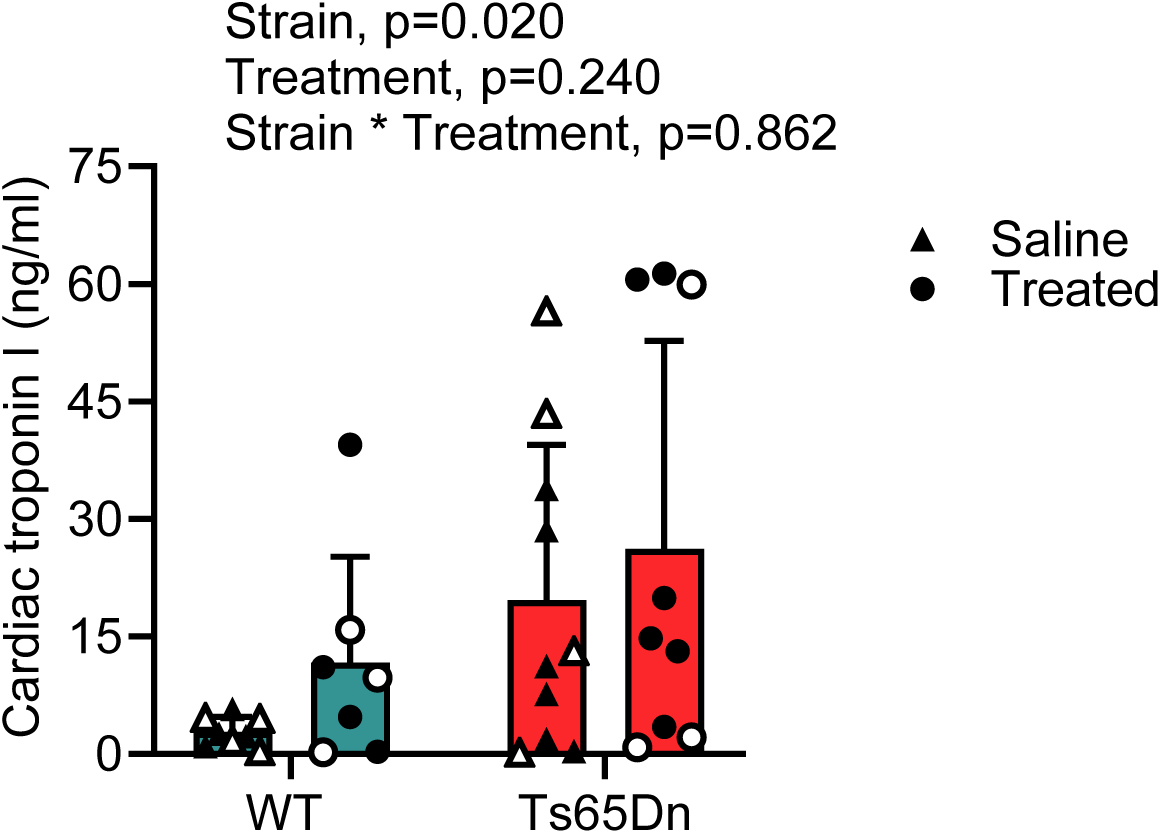
Cardiac troponin. Cardiac troponin I was higher in the serum of Ts65Dn mice (strain, p = 0.020). All data are represented as mean ± SD. Closed shapes represent female mice, and open shapes represent male mice.

## DISCUSSION

The overall purpose of this study was to evaluate the pathophysiology of chronic anthracycline cardiovascular toxicity in the adult Ts65Dn model of Down syndrome. In support of what we know in humans, there was an increased sensitivity of Ts65Dn mice to anthracyclines, reflected by a decrease in the ejection fraction (18). However, the decrease in left ventricular and vascular collagen, and left ventricular mass suggests a unique pattern of cardiac remodeling in Ts65Dn mice.

### Rationale to study anthracyclines in an adult model of DS

This study was motivated by two recent findings in our group in human patients (4, 7). First, adults with DS have an increased risk of cardiovascular diseases, which cannot be predicted by traditional cardiovascular risk factors used for the adult control population, indicating the uniqueness of this group (7). It is a commonly held belief that solid tumor cancers are reduced in adults with Down syndrome, and the risk of blood cancer is the highest in children (19). However, we previously found that adults with DS also have an increased risk of hematological malignancies like leukemia and multiple myeloma, and a similar risk of lymphoma and soft tissue cancers (4). Anthracyclines are part of the standard induction and first-line chemotherapy regimens for several of these cancers (5, 20, 21). Second, adults with DS have an increased overall risk of cardiovascular diseases, including heart failure after anthracycline treatment (4).

The mechanisms of anthracycline toxicity have been extensively studied in the adult control population. It begins with injury to the myocardial cells, leading to a progressive, asymptomatic decline in left ventricular ejection fraction and overt heart failure if left untreated (5). Vascular toxicity is the second most commonly reported cardiovascular toxicity related to anthracycline chemotherapy, and the second most frequent cause of death among patients with cancer receiving these treatments (22).

Since the mechanism of cardiovascular disease development is multifactorial, it is unclear if the mechanism of anthracycline cardiotoxicity in the DS population follows the same pattern, particularly when administered in adulthood. This was confirmed in our study, where we identified a distinct pattern of structural remodeling in Ts65Dn-treated mice compared to WT-treated mice. This supports the hypothesis that although anthracyclines lead to cardiovascular disease in both populations, the structural mechanisms underlying its development differ.

### Daunorubicin toxicity protocol for Ts65Dn mice

Anthracyclines, including daunorubicin, are commonly incorporated in the treatment regimens of patients with DS and myeloid leukemia, which are more effective compared to patients without DS (6, 23, 24). Anthracycline cardiotoxicity is classified into acute and chronic cardiotoxicity (25). Acute toxicity occurs during and up to two weeks after treatment (25). Chronic toxicity, which occurs within and after the first year of treatment, clinically manifests as an asymptomatic decline in left ventricular function that can gradually lead to overt heart failure if left untreated (25, 26).

Unlike in children with DS and acute leukemias, where the recommendation is to reduce the cumulative dose because of the increased sensitivity of DS myeloid leukemia blasts to anthracyclines, there are no dose reduction guidelines for adults (23, 27). We aimed to identify a model that was not acutely toxic but could potentially generate a chronic model of heart failure. Our results showed that matched doses of anthracyclines in Ts65Dn and WT mice led to distinct patterns of cardiovascular remodeling, even with a sublethal dose.

### Implications of collagen reduction

Anthracyclines can reduce left ventricular mass independent of ejection fraction, due to muscle atrophy, cardiomyocyte apoptosis, impaired protein synthesis, and increased oxidative stress (28–31). Anthracyclines can also activate cardiac fibroblasts and induce a metabolic switch toward glycolysis, contributing to fibrotic remodeling (32).

Similar to previous findings in anthracycline cardiotoxicity, we observed an increase in fibrosis in the left ventricles of WT treated mice compared to WT mice treated with saline (33). Remarkably, we observed a decrease in collagen in the left ventricle and thoracic aorta of Ts65Dn-treated mice. The consequences of collagen accumulation in the heart and vessels are relatively well-established (34). However, the effects of decreased collagen on cardiovascular function have been poorly studied (35). Further, the mechanism by which decreased collagen impacts ejection fraction is poorly understood.

There is precedent in genetic models of collagen deficiency, such as osteogenesis imperfecta and Ehlers-Danlos Syndrome, where a reduced ejection fraction is observed (35–38). It is important to note that these syndromes are characterized by defects in collagen types I, III, and V, which play essential roles in maintaining the integrity of the extracellular matrix of the cardiovascular system (39, 40). Therefore, the fact that our results resemble this phenotype suggests a probable similarity in the mechanism of cardiovascular disease remodeling in DS, osteogenesis imperfecta, and Ehlers-Danlos Syndrome. Hence, future physiological research may focus on the importance of collagen underproduction and the mechanism by which it leads to cardiac dysfunction.

Cardiac fibroblasts and vascular smooth muscle cells secrete extracellular matrix components like collagen during injury to support tissue repair and maintain integrity (34). Our findings, which show a decrease in vessel thickness, may be due to the loss of smooth muscle cells, which could potentially weaken the vasculature.

### Comparison of the human DS phenotype and the Ts65Dn mouse model

In our previous study of the incidence of cardiovascular diseases in individuals with DS after chemotherapy, we showed that lifestyle and clinical risk factors of cancer and cardiovascular diseases, such as obesity, tobacco use, and diabetes, were different for adults with DS compared to adults without DS (4). The Ts65Dn mouse model allows us to evaluate the genetic influences of triplicated genes in the absence of confounding lifestyle and clinical variables.

Individuals with DS have low blood pressure and heart rate at rest due to autonomic dysregulation, which is also observed in Ts65Dn mice (10, 41, 42). It is essential to note that although the Ts65Dn mouse model recapitulates many phenotypes of DS, it is trisomic for ∼56% of protein-coding genes on human chromosome 21 (43). Due to the presence of the aberrant piece of mouse chromosome 17 in the Ts65Dn mouse model, results should be interpreted with caution, as some phenotypes may not be biologically relevant to DS (44, 45). Future studies should consider using the recently described Ts66Yah and TcMAC21 mouse models, which were not available at the time of this work but are more closely related to the human DS genotype (46, 47). Also, considering a DS model with lifestyle exposures like obesity could help evaluate how genetics and lifestyle-relevant exposures interact to affect cardiovascular function after anthracycline treatment.

### Genetics of DS

Anthracyclines induce oxidative stress and upregulate extracellular matrix components such as collagen, matrix metalloproteinases, and downstream targets of the transforming growth factor beta (TGF-β) signaling pathway (33). In diseased conditions, collagen is stimulated by factors including chronic activation of TGF-β1, oxidative stress, and persistent inflammation to stabilize damaged tissue after injury (48). Persistent oxidative stress in DS due to the overexpression of genes in the Down syndrome critical region, including *SOD1*, *ETS2*, and *S100,* may disrupt the fibrotic remodeling cascade (49). Excessive accumulation of mitochondrial reactive oxygen species and tumor necrosis factor alpha-mediated inflammation in doxorubicin-treated C57BL/6J mice leads to endothelial dysfunction and arterial stiffness (50, 51).

Previous studies of daunorubicin on rabbits revealed increased apoptosis in the left ventricular myocardium with a corresponding increase in downstream caspases 3 and 7 (52). Hence, in Ts65Dn mice, anthracyclines could induce apoptosis in cardiomyocytes and vascular smooth muscle cells due to elevated oxidative stress and altered metabolism, which may impair their ability to generate a robust fibrotic response and impair ejection fraction. Anthracyclines may preferentially induce cardiomyocyte apoptosis without the ability of the Ts65Dn mice to exhibit a strong fibrotic response, most likely due to excessive oxidative stress and altered metabolism.

The overexpression of trisomy 21-related *RCAN1* and *DYRK1A* inhibits calcineurin, a serine/threonine phosphatase that dephosphorylates the nuclear factor of activated T cells (53). This pathway is essential in the activation of fibroblasts, collagen synthesis, and angiogenesis (54, 55). Angiogenesis has been implicated in the initiation and propagation of fibrosis (56). Therefore, impaired angiogenesis in Ts65Dn mice may contribute to the reduction in collagen, which is critical for myocardial and vascular repair after anthracycline injury.

*Carbonyl reductases* (*CBR*) *1* and *3*, found on human chromosome 21, catalyze the reduction of anthracyclines to cardiotoxic alcohol metabolites (57). Polymorphisms in *CBR1* and *CBR3* influence the synthesis of these metabolites (57, 58). Increased *CBR1* expression and polymorphisms in *CBR3* have been observed in the hearts of donors with DS compared to donors without DS, which drives the cardiac synthesis of daunorubicinol (58). Therefore, daunorubicinol may accumulate in the heart and vessels of Ts65Dn mice, contributing to cardiomyocyte loss and impaired fibrotic signaling. Also, polymorphisms in *CBR3* have been associated with a decline in left ventricular ejection fraction after anthracycline therapy (59, 60). Additionally, the overexpression of interferon receptor genes on human chromosome 21 contributes to immune dysregulation in DS (61, 62). Persistent interferon signaling can suppress fibrosis (63).

Doxorubicin increases the transcription of histone deacetylases and decreases DNA methylation in the heart of male Wistar-Han rats (64). While recent studies have focused on the possibilities of circulating microRNAs as biomarkers for anthracycline cardiotoxicity, it has not been studied in the context of DS (65). Transcriptional changes have been observed in DS phenotypes, which have been associated with the gene dosage imbalance on human chromosome 21, but most genes encoding microRNAs and long non-coding RNAs are not mapped to human chromosome 21 (66, 67). Future studies should investigate the impact of anthracycline exposure on gene expression in this model, including non-coding gene products.

C57BL/6J mice treated with a low dose of doxorubicin exhibit increased pulse wave velocity four days after treatment and four weeks after a single dose of doxorubicin (50, 68). However, individuals with DS exhibit no significant difference in aortic stiffness compared to age- and sex-matched controls without DS, a phenomenon also observed in Ts65Dn mice (10, 69). This aligns with our findings of no change in pulse wave velocity between WT and Ts65Dn mice. The increased expression of genes on human chromosome 21 involved in cardiovascular function can create an environment that modifies treatment response in DS. Although direct evidence for this interaction in DS is limited, our findings suggest that the increased expression of these genes may alter chemotherapy outcomes, highlighting the need for integrated cardio-oncology assessments in this population. This relationship may provide the basis for the likelihood of a ‘reverse cardio-oncology’ phenomenon in this population. Reverse cardio-oncology is an emerging concept in cardio-oncology, which suggests that underlying cardiovascular diseases can increase cancer risk and influence cancer outcomes (70).

### Cardiac troponin as a biomarker for DS in anthracycline cardiotoxicity

The leakage of cardiac-specific troponins I and T into the circulation indicates early myocardial injury and are biomarkers of acute anthracycline cardiotoxicity (57). We found that cardiac troponin I did not increase in the wild-type model, indicating we had surpassed the acute phase of cardiac damage. Increased *DYRK1A* expression is associated with an increase in the expression of fetal cardiac troponin T variants in myocardial tissues of donors with and without DS (71). Cardiac troponins I and T are expressed at similar levels in the myocardium (72). Therefore, the increased expression of *DYRK1A* that results from gene triplication may also lead to an increase in cardiac troponin I expression in the Ts65Dn mice.

Given the elevated cardiac troponin I we observed, even at baseline, it raises concerns about whether it is an effective marker of myocardial injury for the DS population and highlights the need to investigate additional biomarkers or define DS-specific normative values. Additional studies on baseline troponin levels in the DS population are needed to determine a reference value of cardiac troponin for DS and to validate the utility of cardiac troponin as a biomarker for anthracycline cardiotoxicity in this population.

### Limitations

We had the advantage of evaluating cardiac function by intracardiac catheterization, which, by necessity, is a terminal procedure. However, future studies should consider serial monitoring of cardiac function to identify progressive changes in ejection fraction. We have previously shown that urethane anesthesia significantly elevates heart rate and depresses arterial oxygen saturation and blood pressure in C57BL/6J mice (15). Autonomic dysregulation by urethane anesthesia might have influenced cardiovascular outcomes from the left ventricular catheterization, and future studies should quantify its impact in DS-relevant models. Since non-coding genes have been underexplored in DS, most of our interpretations were based on the coding genes. The Ts65Dn mouse model of DS contains other triplicated genes that may be unrelated to the DS phenotype (44). Other newly developed mouse models, such as the Ts66Yah and TcMAC21 models, do not have the triplication of DS-unrelated genes (46, 47). Even though the cardiovascular phenotypes of these models have not been extensively studied, it is worth replicating our findings in these models. Additionally, we dosed mice at a single developmental time point. Due to the increasing life expectancy of individuals with DS, it is important to consider the effects of anthracyclines when administered at multiple ages. Finally, arterial stiffness is pressure-dependent, and therefore, an increase in blood pressure leads to an increase in pulse wave velocity (73). Future studies should consider normalizing pulse wave velocity with blood pressure to account for its functional effects on arterial stiffness.

### Conclusions

We have shown a phenotype that recapitulates what is observed in humans: the Ts65Dn mouse model of Down syndrome is more sensitive to anthracyclines, but the pathway to a reduced ejection fraction may be different. The deviation from the established pathogenesis of anthracycline cardiotoxicity underscores the need for more DS-related mechanistic studies to refine DS-specific cancer treatment recommendations.

## DATA AVAILABILITY

Data will be made available upon reasonable request to the corresponding authors.

## SUPPLEMENTAL MATERIAL

Supplemental Tables S1-S3.

## ACKNOWLEDGMENTS

The authors acknowledge Chantal Allamargot, PhD; David Gordon, MD, PhD; Jessica Gorzelitz, PhD; Jordan Turner; Lara DeRuisseau, PhD, and Zishan Zhang, PhD, for their valuable feedback on this work.

## Author contributions

MAB, AV, MHT, MLB – Conceived and designed research; MAB, AV – performed experiments; MAB, AV, HK, IE, CRJ, MHT, MLB – analyzed data; MAB, AV, HK, IE, CRJ, BL, GB, PB, MHT, MLB – interpreted results of experiments; MAB, AV, MHT, MLB – drafted manuscript; MAB, AV, HK, IE, CRJ, GM, MV, BL, GB, PB, LRD, MHT, MLB – edited and revised manuscript; MAB, AV, HK, IE, CRJ, GM, MV, BL, GB, PB, LRD, MHT, MLB – approved final version of the manuscript.

## SUPPLEMENTAL TABLES

**Supplemental Table 1.** Survival curve comparisons.

| Curve Comparison | p-value |
| --- | --- |
| WT saline vs WT 2mg/kg | 0.317 |
| WT saline vs WT 4mg/kg | 0.027 |
| Ts65Dn saline vs Ts65Dn 2mg/kg | 0.070 |
| Ts65Dn saline vs Ts65Dn 4mg/kg | **<0.001** |

All statistically significant values are indicated in bold. The Bonferroni correction was used for multiple comparisons, $p < 0.006$ .

**Supplemental Table 2.**
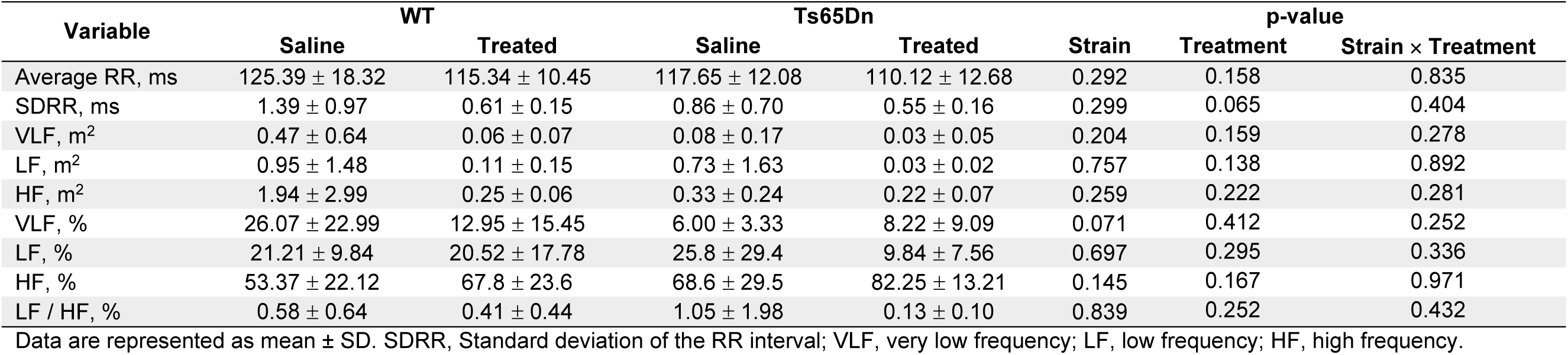
Summary statistics for heart rate variability.

**Supplemental Table 3.**
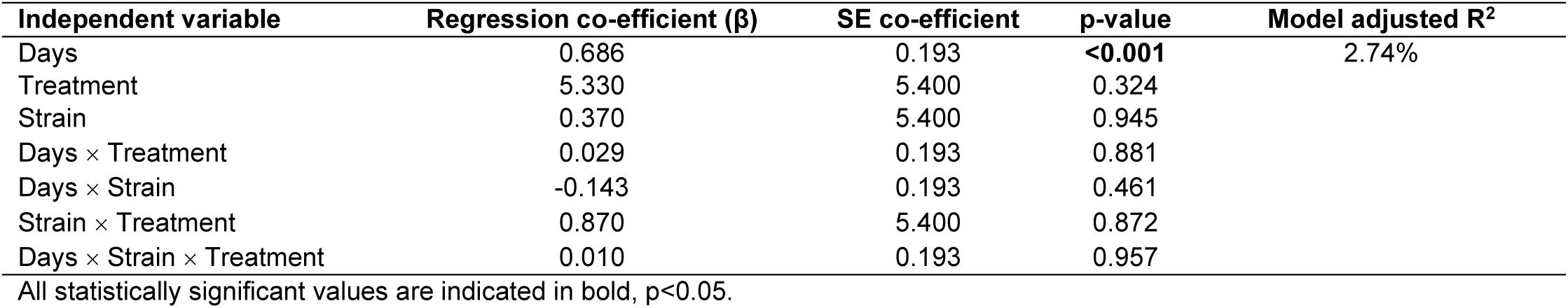
Analysis of co-variance statistics of pulse wave velocity measurements.

| Independent variable | Regression co-efficient (β) | SE co-efficient | p-value | Model adjusted R<sup>2</sup> |
| --- | --- | --- | --- | --- |
| Days | 0.686 | 0.193 | **<0.001** | 2.74% |
| Treatment | 5.330 | 5.400 | 0.324 |  |
| Strain | 0.370 | 5.400 | 0.945 |  |
| Days × Treatment | 0.029 | 0.193 | 0.881 |  |
| Days × Strain | -0.143 | 0.193 | 0.461 |  |
| Strain × Treatment | 0.870 | 5.400 | 0.872 |  |
| Days × Strain × Treatment | 0.010 | 0.193 | 0.957 |  |

All statistically significant values are indicated in bold, $p < 0.05$ .

